# Cell-cultured PDMS vascular model to allow placement of implant devices

**DOI:** 10.1101/2025.01.20.634010

**Authors:** Taku Okuno, Kazuyo Ito, Kenichi Funamoto, Daisuke Yoshino

## Abstract

**Background:** A stent maintains normal blood flow by expanding a stenotic artery from the inside. Stents currently in clinical use are mechanically sub-optimized and, therefore, exert excessive mechanical stimuli on the vascular wall. This can lead to arterial inflammation and potentially in-stent restenosis. To optimize the force exerted by a stent on the vascular wall, we should understand, reproduce, and analyze in detail the mechanical field to which the stented vessel is exposed. However, to date, no *in vitro* model has been established that can adequately evaluate the mechanical effects of a stent on the vascular wall and be constructed without relying on a laboratory’s know-how. This study aims to develop a three-dimensional *in vitro* vascular model that allows stent placement and to provide the details of its construction method.

**Methods:** Polydimethylsiloxane (PDMS) was used to mimic the adventitial structure of arteries. Human carotid artery endothelial cells (HCtAECs) were then seeded on the luminal surface and cultured for 24 h to form a confluent monolayer (intima). The constructed model was installed in the flow-exposure culturing system, and hemodynamic stimulation (two types of shear stress (SS); 0.5 Pa and 2.3 Pa) was applied to the HCtAECs inside to reproduce the physiological state of blood vessels. In addition, a self-expanding stent was placed in the model during perfusion culture.

**Results:** We examined the performance of the developed model based on quantitative evaluations of endothelial morphology in response to SS and in-stent endothelialization. Exposure to SS for 24 and 48 h caused endothelial orientation and elongation in the direction of flow, confirming the physiological responses of blood vessels. We were also able to quantitatively evaluate endothelial migration (endothelialization) to the stent.

**Discussion:** Our model can reproduce the mechanical field inside the stented blood vessel, including hemodynamic stimuli. Based on the results obtained, the developed cell-cultured vascular model has sufficient performance to be used not only for quantitative evaluation of in-stent endothelialization but also for optimization of the mechanical fields in the stented vessel.

## INTRODUCTION

Stent placement has been widely used as a minimally invasive treatment for vascular stenosis associated with atherosclerosis of the carotid or coronary arteries. Stents have a mesh-like cylindrical structure and ensure blood flow through stenotic lesions by expanding the vascular lumen like a spring. Most stents currently in clinical use have been modified with drugs to prevent restenosis (*i.e.*, drug-eluting stents). Meanwhile, the mismatch between stent stiffness and vessel stiffness remains unresolved, making it difficult to suppress the inflammatory response of the vascular wall to the excessive mechanical stimulation caused by the stent placement, and the risk of late thrombosis and restenosis tends to be high [1–4]. Optimizing the mechanical stimulation of the stent on the vessel wall, which is one of the main causes of vascular inflammation, is necessary to maximize the advantages of stent placement as a minimally invasive treatment. However, the effect of the mechanical stimulation has not been studied in detail, even though it is much greater than the shear stress generated by blood flow [5–8]. Consequently, the strategy to optimize the stiffness of the stent remains unclear.

To optimize the force exerted by a stent on the vascular wall, we should first understand the mechanical field to which the stented vessel is exposed. The most useful information can be obtained from stent placement in humans [9,10] or experimental animals [11,12] (*i.e.*, *in vivo* trials), but in addition to large individual differences, ethical issues limit the quantitative evaluation of the mechanical effects of stents on the blood vessel wall. In the field of tissue engineering, *in vitro* vascular construction by means of bioprinting and microfluidic device technology is currently undergoing rapid development [13,14]. However, these models are primarily focusing on microvascular networks that deliver nutrients and oxygen to tissues or organoids [15–17], and there are few models with vessel diameters large enough to allow stent placement. Some studies have actually placed stent struts on a surface culturing vascular endothelial cells (ECs) and evaluated their endothelialization [18–20], but they do not reproduce the mechanical field to which stents and blood vessels are exposed. Although those studies can be used to examine the biocompatibility of surface treatments (including chemical modification and micro-grating), they are not suitable for optimizing the mechanical forces exerted by stents on the vascular wall. On the other hand, a perfusion culture system using a small-to-medium vascular model with cultured endothelial cells has been developed to allow stent placement, enabling us to analyze the cellular dynamics of stented blood vessels [21–23]. The previous studies have adopted models with an outer membrane made of polydimethylsiloxane (PDMS) or fluororesin, and some of them have a collagen medium with vascular smooth muscle cells (SMCs) inside. They have enabled detailed analysis of EC dynamics in a stented blood vessel. However, our knowledge of how to optimize the mechanical force exerted by the stent on the vessel wall is still limited. In addition, the method to construct the vascular model is complicated and requires lots of know-how, making it difficult for others to reproduce.

We have worked to develop a design theory for a stent that can regulate its stiffness in a position-dependent manner [24, 25], and to construct a culture system that provides spatiotemporal control of hemodynamic stimuli (shear stress [26, 27] and hydrostatic pressure [28, 29]). Here, we construct a cell-cultured PDMS vascular model that allows stent placement, following the previous studies, and install it in our flow exposure system [26, 27]. We design the construction method to be as simple as possible and demonstrate the details. This study also proposes its possibility as a general application for stent evaluation by verifying the reproduction of vascular dynamics and in-stent endothelialization. We expect that the combination of this technique and our design theory [24, 25] will yield to quantify the relationship between the mechanical forces that stents exert on the vascular wall, and restenosis or neointimal hyperplasia, and to determine the optimal conditions for stent stiffness, which is an open problem.

## MATERIALS AND METHODS

### Cell culture

Human carotid artery ECs (HCtAECs; lot no. 2463, 3014-05a, Cell Applications, San Diego, CA, USA) were cultured in Medium 199 (M199; 31100-035, Gibco, Thermo Fisher Scientific, Waltham, MA, USA) containing 20% heat-inactivated fetal bovine serum (FBS; 173012, Sigma-Aldrich, St, Louis, MO, USA), 10 µg/L human basic fibroblast growth factor (bFGF; GF-030-3, Austral Biologicals, Ango, San Ramon, CA, USA), and 1% penicillin/streptomycin (P/S; 15140-122, Gibco). HCtAECs from the fifth to ninth passages were used for experiments in this study.

### Fabrication of a PDMS vascular model

A straight channel vascular model with a circular cross-section was fabricated from PDMS (Sylgard 184 Silicone Elastomer Kit, Dow Chemical, Midland, MI, USA). Connectors, which serve as the inlet and outlet of the vascular model, were first prepared by casting and curing PDMS in polyacetal molds (**Fig. 1a**). A stainless-steel core rod was then inserted through the connectors and sandwiched between the PDMS-filled upper and lower molds, which form the straight flow channel of the vascular model, and cured (**Fig. 1b**). A degassing procedure was performed after the PDMS was poured into the mold to remove air bubbles from entering the vascular model. Finally, three types of PDMS vascular models with inner and outer diameters of 4 mm and 5 mm, 5 mm and 6 mm, or 6 mm and 7 mm, respectively, were constructed by immersing them in anhydrous ethanol and removing the core rods (**Fig. 1c**).

**Fig. 1.**
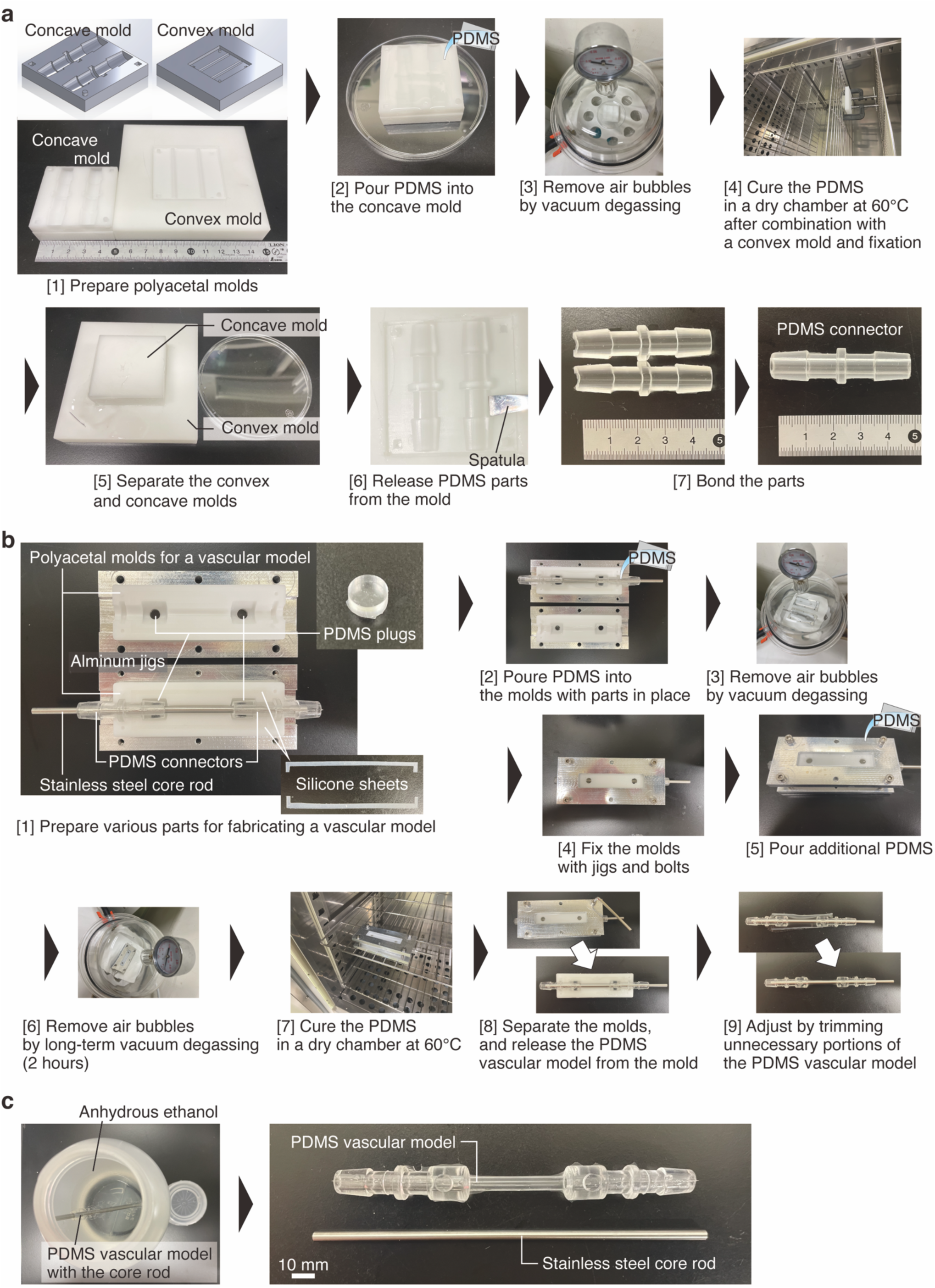
Fabrication of a PDMS vascular model. (**a**) Procedures for preparing connectors that form the inlet and outlet of the vascular model. (**b**) Steps for constructing the model. (**c**) Instructions on how to remove the core rod from the model.

### Construction of a cell-cultured vascular model

A cell-cultured PDMS vascular model was constructed by seeding and forming a monolayer of HCtAECs on the lumen of the fabricated vascular model with an inner diameter of 4 mm and an outer diameter of 5 mm. L-shaped fittings (AJL10, ISIS, Osaka, Japan), silicone tubes (96400-18, Masterflex, Cole-Parmer, Barrington, IL, USA), and the PDMS vascular model were washed with ultrapure water and autoclaved (**Fig. 2a**). After drying for 24 h, they were hydrophilized with a plasma cleaner (PDC-32G, Harrick Plasma, Ithaca, NY, USA) for 3 min, and immediately assembled. Following that, 2.5 mL of a 4:1 mixture of 0.2% gelatin solution (G9391, Sigma-Aldrich) and poly-D-lysine hydrobromide solution (1 mg/mL; P7886, Sigma-Aldrich) was injected into the lumen of the assembled model. The models were then sealed and placed in a CO_2_ incubator and incubated for 4 h, changing the circumferential angle every hour in the order of 0°, 90°, 180°, and 270° (**Fig. 2b**). After incubation, the solution was drained, and the models were dried at 60°C for 24 h. After washing the lumen of the models with phosphate-buffered saline (PBS; 05913, Shimadzu Diagnostics Corp., Tokyo, Japan), a suspension of HCtAECs prepared at a concentration of 5.0×10^5^ cells/mL was injected and incubated for 7 min 30 s and 22 min 30 s at a circumferential angle of the PDMS vascular model of 0° and 30°, respectively. Similarly, the suspension was re-injected and incubated for 7 min 30 s and 22 min 30 s at the tilt angle of 180° and 210°, respectively (**Fig. 2c**). An endothelial monolayer was then formed by slowly changing the culture medium and incubating for 24 h to prevent cell detachment.

**Fig. 2.**
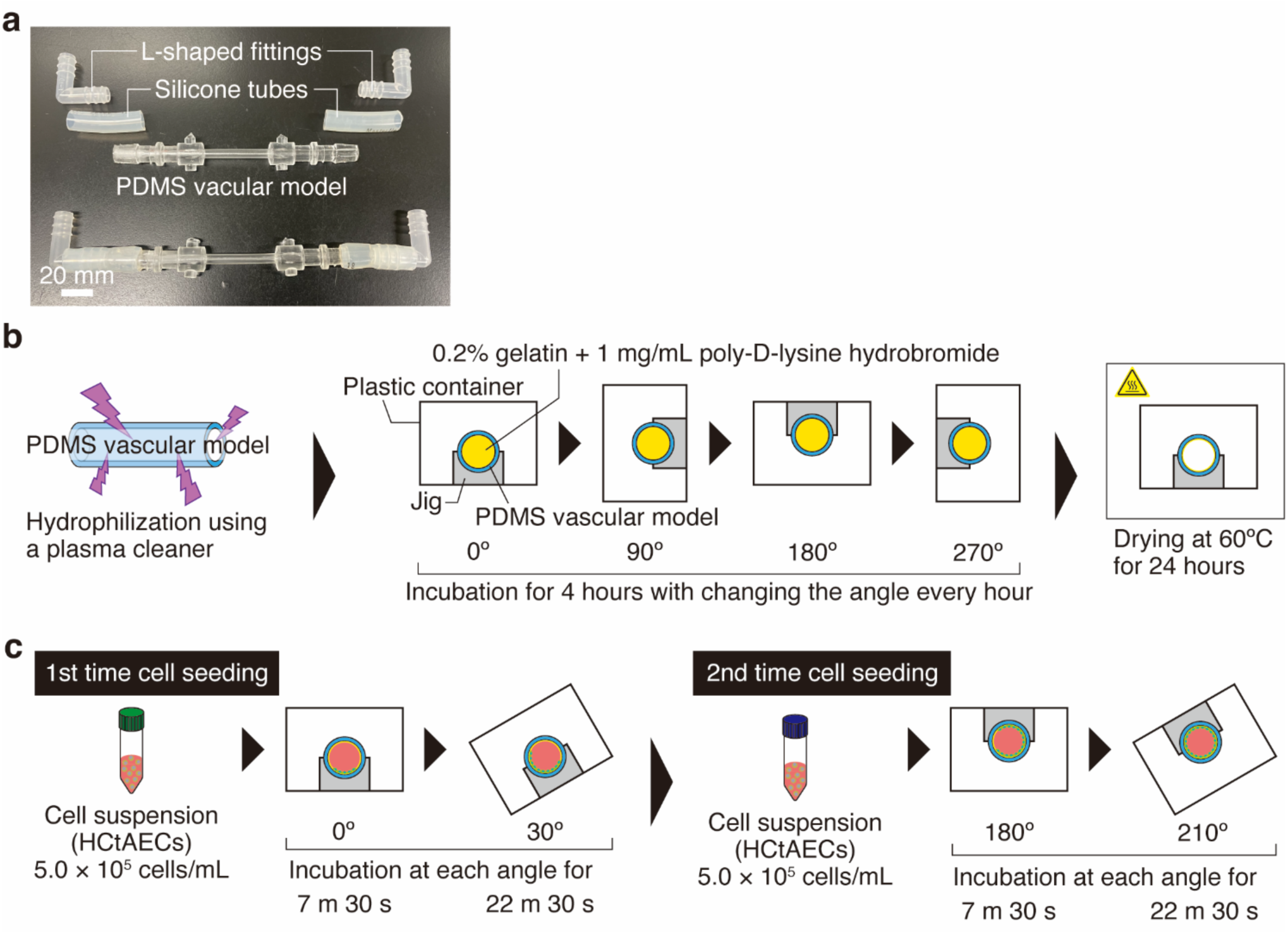
Construction of a cell-cultured PDMS vascular model. (**a**) Configuration of the vascular model. (**b**) Hydrophilization treatment of the model and coating with extracellular matrix (gelatin and poly-D-lysine). (**c**) Protocol for cell seeding into the model lumen.

### Exposure to shear stress

To mimic vascular physiology, HCtAECs were exposed to shear flow by connecting the constructed model to a flow circuit consisting of a peristaltic pump (7528-10, Masterflex, Cole-Parmer) and two reservoirs (**Supplementary Fig. 1a**) [26, 27]. The flow circuit was filled with M199 with Hanks’ balanced salt (M0393, Sigma-Aldrich) supplemented with FBS, bFGF, and P/S at the same concentrations as the culture medium to maintain the pH at 7.4 throughout the experiment. The circuit was also maintained at 37°C using a temperature-controlled water bath. Cells were exposed to two different fluid shear stresses (SS) for up to 48 h, whereas those incubated in a static condition without SS were defined as “static.” The flow rate was set to 4.19 or 8.38 mL/s.

### Wall shear stress estimation

The two flow rates correspond to Reynolds number of 1,778 (4.19 mL/s) and 3,557 (8.38 mL/s), respectively, in the PDMS vascular model. The SS acting on the model wall was determined by the following equation about SS (*τ*) on a circular pipe wall:

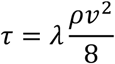

where *λ*, *ρ*, and *ν* represent the pipe friction coefficient, the density of the fluid, and the average flow velocity, respectively. As for the former condition of 4.19 mL/s, the flow to of culture medium was estimated to be laminar flow, and therefore the pipe friction coefficient was obtained as follows:

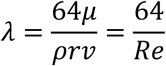

where *μ*, *r*, and *Re* represent the viscosity of the fluid, the radius of the circular pipe, and the Reynolds number, respectively. The viscosity *μ* and density *ρ* of the culture medium were 0.75 mPa·s and 1,000 kg/m^3^, respectively. The radius of the tube *r* was 2.0 mm. On the other hand, as for the latter condition, the flow could be disturbed because the Reynolds number was >2,300. Hence, the pipe friction coefficient was obtained based on the Blasius equation for the friction factor for turbulent flow in a smooth pipe:

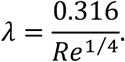

By substituting the parameters into the equations, the SS acting on the model wall was obtained as 0.5 and 2.3 Pa under the slow and fast flow, respectively.

### Stent placement into the cultured vascular model

The constructed cultured vessel model can be placed with a stent-like implant device. In this study, we observed cellular dynamics in the model implanted with a self-expanding stent made from nickel-titanium (NiTi) alloy. The stent with an outer diameter of 6 mm (SENDAI stent [24, 25]) was first shrunk to approximately 4 mm by cooling in water with ice, and then inserted into an industrial straw with an inner diameter of 2.6 mm (KP030026, SHIBASE, Okayama, Japan) by applying further shrinkage using a tapered pipe (**Fig. 3a**). Subsequently, the stent mounted in the straw was sterilized by immersion in 70% ethanol. After 24-h application of shear flow in the cultured vascular model in which the L-shaped fitting on the upstream side had been replaced with a T-shaped fitting (AJT10, ISIS; **Supplementary Fig. 1b**), the flow was stopped, and the silicone tubes on the upstream and downstream sides were tied respectively to completely stop the flow of culture medium (**Fig. 3b**). A stent mounted in the straw that had been rinsed with sterile ultrapure water and PBS was inserted from the T-shaped fitting, pushed out using a sterile rod, and left in place in the cultured vascular model (**Fig. 3c**). After the stent was placed, shear flow was reapplied, and the cells were cultured for up to 48 h.

**Fig. 3.**
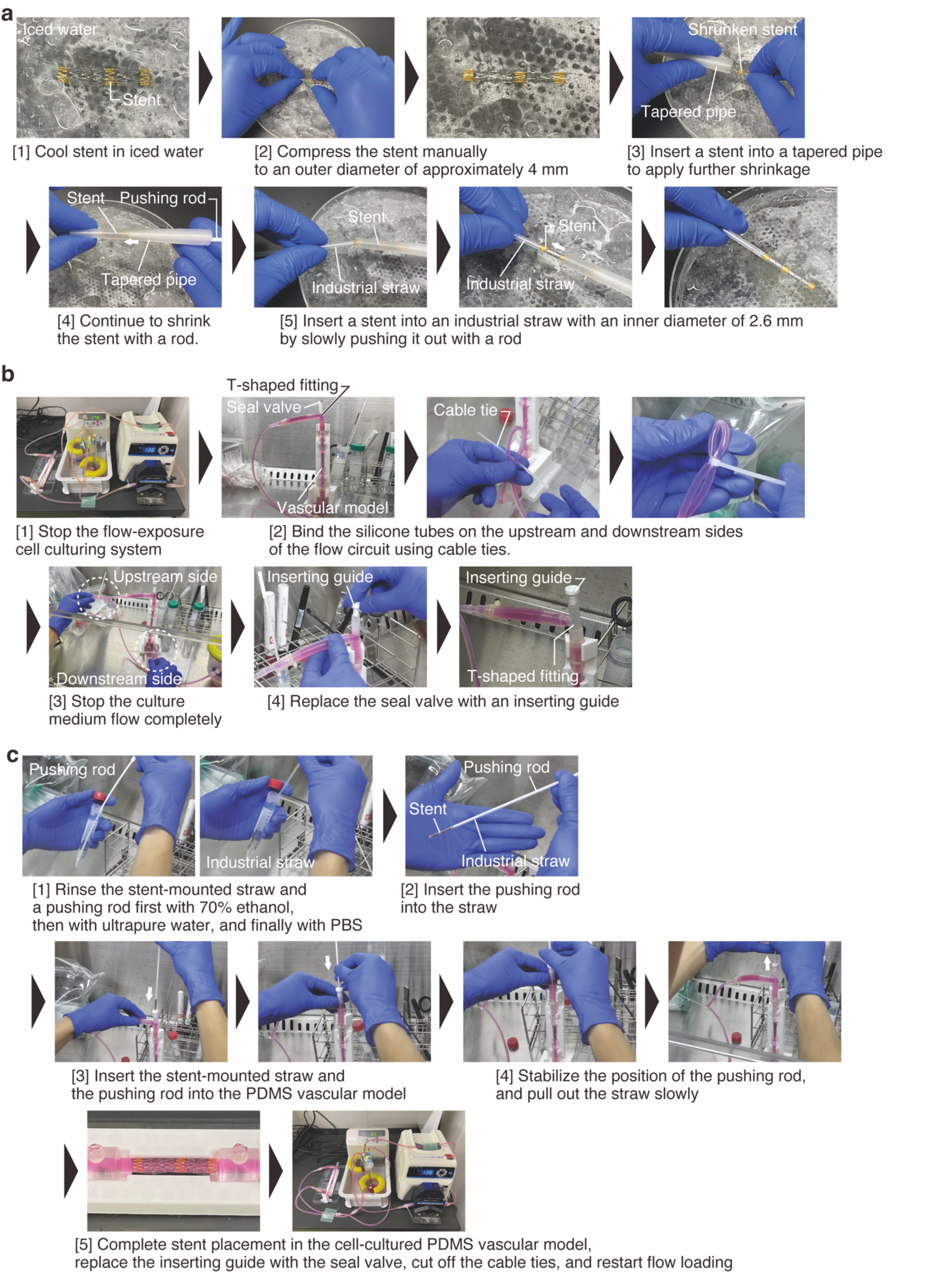
Procedures for stent placement into the cell-cultured PDMS vascular model. (**a**) The processes of shrinking a stent and mounting it on an industrial straw. (**b**) Preparation for stent placement: How to temporarily stop the flow-exposure experiment. (**c**) Instructions for inserting the stent into the vascular model.

### Immunofluorescence staining

After exposure to SS, ECs were fixed with 4% paraformaldehyde (PFA; 158126, Sigma-Aldrich) in PBS for 10 min at 20–25°C. The fixed ECs were permeabilized with 0.1% TritonX-100 (17-1315-01, Pharmacia Biotech, Uppsala, Sweden) in PBS for 5 min, followed by incubation in PBS containing 1% Block Ace (BA; UKB40, KAC, Kyoto, Japan) for 40 min to prevent nonspecific antibody adsorption. Cells were then stained with primary and secondary antibodies diluted in 1% BA in PBS and PBS (primary antibody; 1:200, secondary antibody; 1:200), respectively. In this study, we used the mouse monoclonal anti-VE-cadherin antibody (sc-9989, Santa Cruz Biotechnology, Dallas, TX, USA) as the primary antibody and Alexa Fluor 594 conjugated goat anti-mouse IgG antibody (A-11005, Thermo Fisher Scientific) as the secondary antibody. Actin filaments and cell nuclei were stained using Alexa Fluor 488 Phalloidin (A12379, Thermo Fisher Scientific) and 4’,6-diamidino-2-phenylindole, dihydrochloride (D1306, Thermo Fisher Scientific), respectively. Alexa Fluor 488 conjugated goat anti-mouse IgG antibody (A-11001, Thermo Fisher Scientific) and Alexa Fluor 594 Phalloidin (A12381, Thermo Fisher Scientific) were used for staining as needed.

### Measurement of fabricated model dimensions

To investigate accuracy and precision in the fabrication of the PDMS vascular model, we measured its inner diameter and wall thickness as reference dimensions. Each model was cut at five locations, and the cross-sections were captured using a stereomicroscope (Stemi 305 cam, Carl Zeiss, Oberkochen, Germany). Based on the obtained microscopic images, the inner diameter of the model was calculated from the lumen area, and the wall thickness was measured at 12 points per section (every 30 degrees). This study fabricated three samples of each diameter and used them for evaluation.

### Morphological evaluation

To evaluate cell morphology, we quantified EC elongation and orientation in the flow direction by means of an aspect ratio and orientation angle. We first extracted the outlines of ECs along an adherens junction protein, VE-cadherin, in the fluorescence images. Based on the extracted outlines, ellipses equivalent to the shapes of the ECs were calculated using ImageJ Fiji [30]. The aspect ratio was defined as the ratio of the minor axis to the major axis of an ellipse equivalent to the shape of the cell. A perfect circle has a maximum aspect ratio of 1, whereas for highly elongated shapes, the aspect ratio approaches zero. The angle between the major axis of the ellipse and the direction of flow was measured as the orientation angle of the cell. At least three independent experiments were conducted, and 50 cells were randomly extracted for each.

### Calculation of detachment rate

To analyze cell detachment, we defined the area of cell detachment relative to the total area of the luminal surface of the vascular model in the microscopic images as the detachment rate. The fluorescence images of F-actin in the cells were first converted to 8-bit images and then binarized according to Otsu’s method [31]. Thresholds were adjusted by comparing the actual microscopic images in cases where the positions of the cells were clearly missing during binarization. Then, the black and white of the image was inverted, and the region where the cells were detached was converted to white. The morphological opening was applied to remove small objects, such as noise, from the binarized image, followed by calculation of the area of the cell detachment region using the ‘Analyze Particles’ function of ImageJ. At least 10 images for each experiment were used in the calculation of detachment rate.

### Quantification of in-stent endothelialization

To quantify in-stent endothelialization, the percentage of cell migration and adhesion to the lateral and inner top sides of the stent was determined as the endothelialization rate. We first painted the side of the stent (either the lateral or top sides) to be analyzed in the fluorescent image in white; then, the non-painted areas were erased, and the image was inverted to black and white (**Supplementary Fig. 2**). Using the ‘Z-Projection (Projection Type: Max Intensity)’ function in ImageJ, the fluorescent staining image of F-actin was superimposed on the target region. Binarization was then performed with a threshold adjustment relative to the microscopic image, and the non-endothelialized area was left white. The area of the non-endothelialized region was calculated and subtracted from the area of the stent side to be analyzed to obtain the area of the in-stent endothelialized region. We analyzed five locations for the top and lateral sides, respectively, in each experiment.

### Statistical analysis

Each data was obtained from at least three independently repeated experiments. Statistical significance was calculated using the two-sided Welch’s *t*-test with Bonferroni correction for multiple comparisons, with statistical significance set at *P* < 0.01 (significant difference). The effect size of each statistical test for each parameter *X* was analyzed using the Cohen’s *d* [32], which is defined as follows:

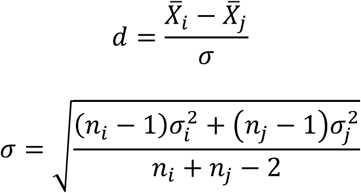

Here, 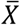, *σ*, *n* are the mean value, standard deviation, and size of sample, respectively. The subscripts *i* and *j* indicate the conditions that are the subject of the comparison test. The exact *P*-values and the effect size for all statistically tested data are described in **Supplementary Data 1**.

The kurtosis (*KURT*) and skewness (*SKEW*) were used as indicators to compare the distribution of EC orientation in the flow direction. These indicators are defined by the following equations [33].

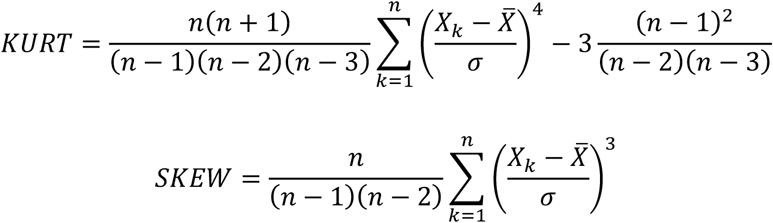

## RESULTS

### Construction of a cultured vascular model

We first investigated the accuracy and precision in the fabrication of PDMS vascular models of three different diameters (**Fig. 4a**). The models were cut at five locations, and their inner diameters and wall thicknesses were measured (**Fig. 4b**). The deviation from the target value for the inner diameter of each model was within ±100 µm, and the measurements were consistently smaller than the target value in the models with inner diameters of 4 mm and 6 mm (**Fig. 4c**). In terms of fabrication precision, the 4-mm inner-diameter models had the smallest standard deviation, followed by the 6-mm models, and the 5-mm models had the largest standard deviation. For wall thickness, the deviation from the target value varied depending on each model, with the 4-mm inner-diameter models being within ±150 µm, the 5-mm models being within ±250 µm, and the 6-mm models being within ±200 µm (**Fig. 1d**). The fabrication precision of the wall thickness followed a similar trend as the inner diameter. We confirmed that it is possible to fabricate PDMS vascular models with high yield using the proposed method (**Fig. 1**), as the dimensions (inner diameter and wall thickness) of all fabricated models were generally within 3σ (three times the standard deviation).

**Fig. 4.**
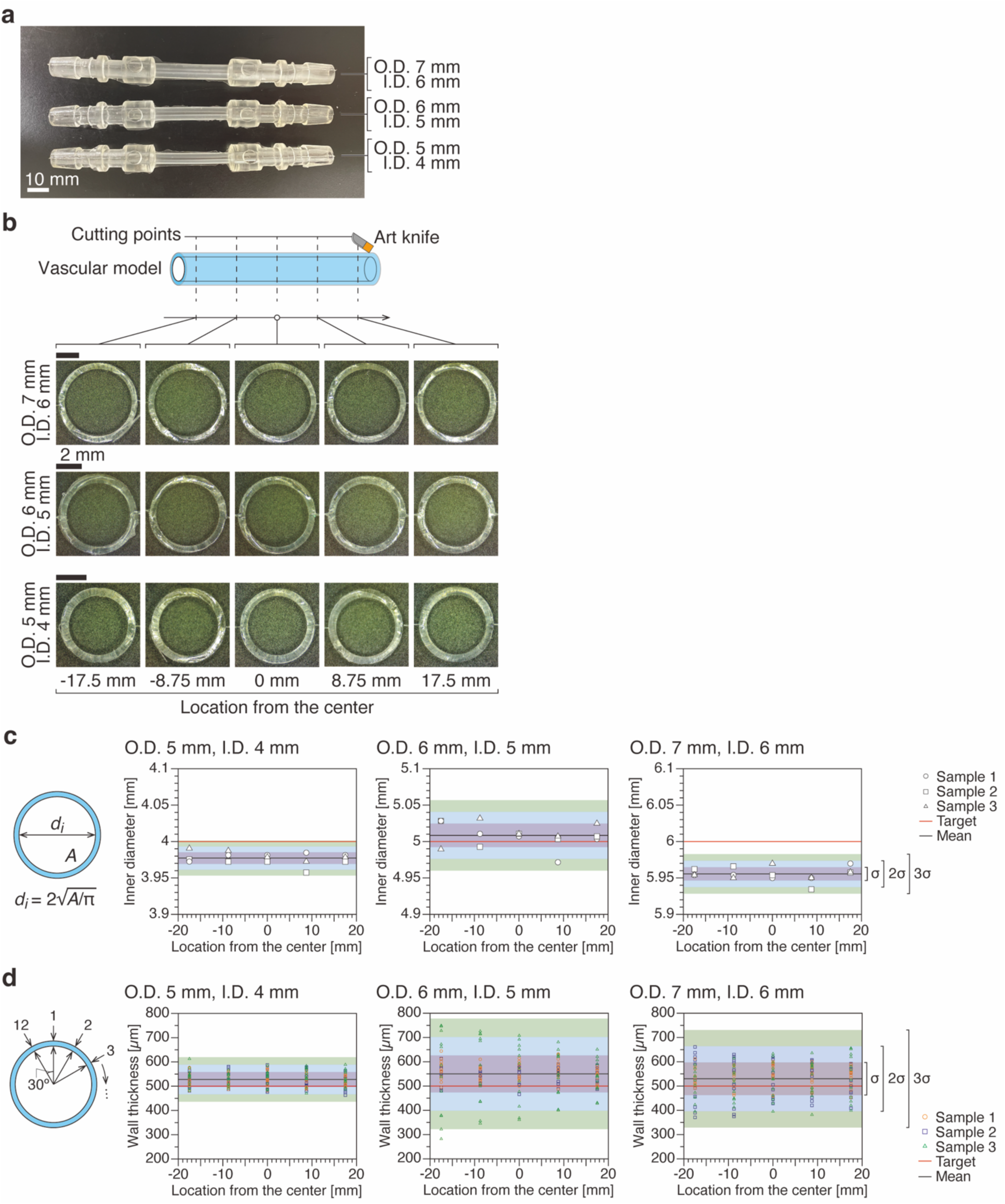
Productivity evaluation of PDMS vascular models. (**a**) Photographs of three types of the fabricated PDMS vascular models. (**b**) Sample segment at each position of the model. Scale bars, 2 mm. (**c**) Measured inner diameter of each model (*n* = 3). (**d**) Measured wall thickness for each model (*n* = 3). Red line: target inner diameter, black line: mean inner diameter of the model. Purple, light blue, and green highlights represent the range of 1x, 2x, and 3x standard deviation (σ), respectively.

We then seeded HCtAECs into a PDMS model of 4 mm inner diameter vessels to construct a cultured vascular model. At 24 h after seeding, the cells covered the model lumen with their monolayer (**Fig. 5a**). In the magnified view, VE-cadherin was observed in the intercellular adhesion sites, and the integrity of the endothelium was maintained without gaps. In the cross-sectional view, the cells were attached in a circumferential direction, forming a cylindrical lumen (**Fig. 5b**). However, in the 48 h after seeding, most of the cells had detached, and they were unable to maintain the monolayer (**Fig. 5c**).

**Fig. 5.**
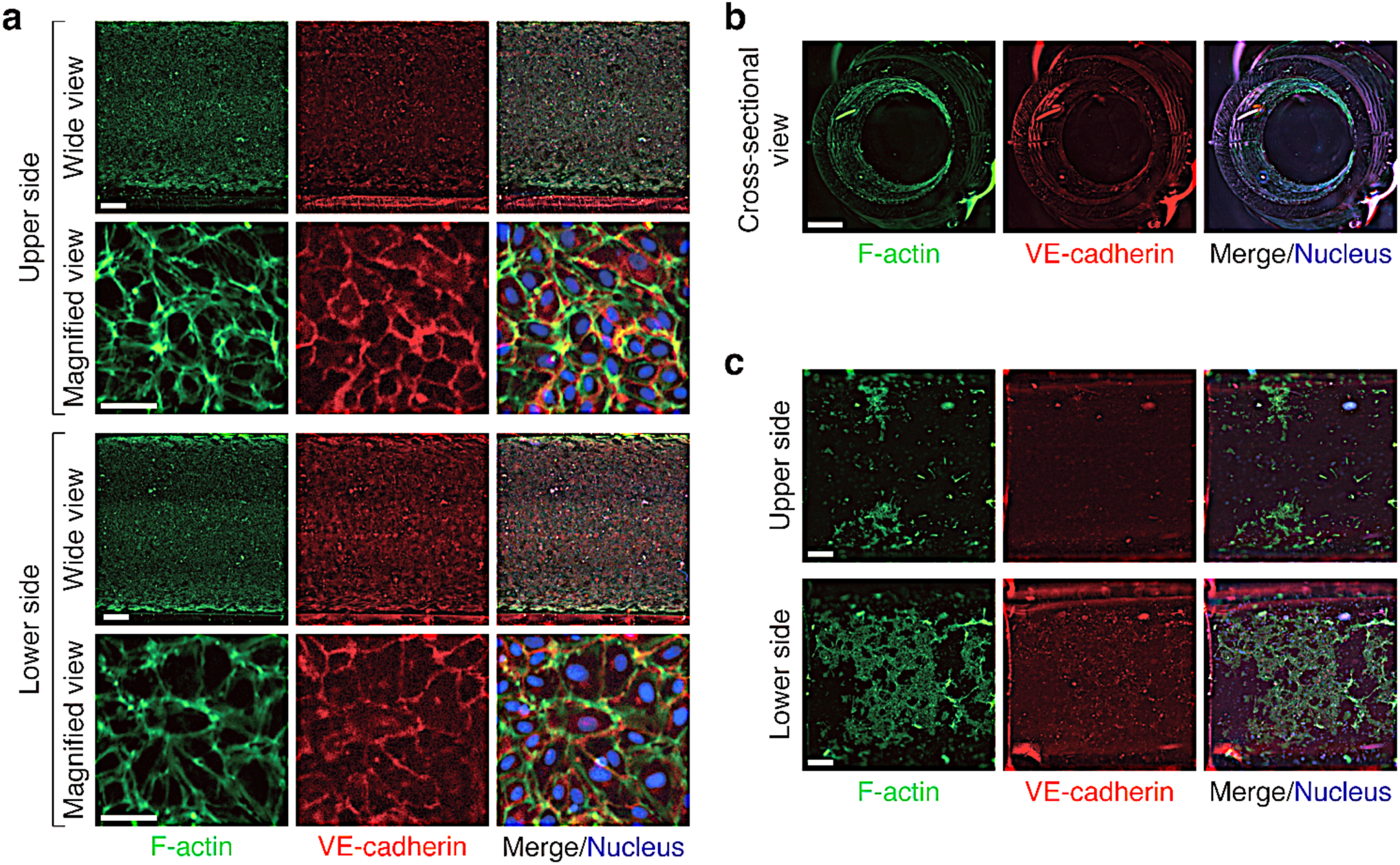
Characteristics of the cell-cultured PDMS vascular models. (**a**) Representative images of fluorescent-stained HCtAECs on the upper and lower sides of the vascular model after 24 h of seeding. Scale bars, 500 µm (wide view) and 50 µm (magnified view). (**b**) Representative images of fluorescent-stained ECs in the cross-section of the model after 24 h of seeding. Scale bar, 1 mm. (**c**) Wide-angle images of fluorescent-stained ECs after 48 h of seeding. Scale bars, 500 µm

### Endothelial response to shear stress in the cultured vascular model

To further examine the characteristics of the cultured vascular model, we connected it to a flow circuit (**Supplementary Fig. 1**) and applied SS to the HCtAECs inside. After 24 h of seeding, we started to apply flow to the cells and analyzed the morphological response of the cells in the 24-h and 48-h exposures. We found no difference in the response of ECs to shear flow on the upper and lower sides. ECs exhibited a polygonal shape under the static condition, whereas they elongated and oriented in the flow direction under SS conditions (**Fig. 6a**). In addition, the development of actin stress fibers parallel to the flow direction was observed at the cell periphery. The expression of VE-cadherin at the intercellular adhesion sites was somewhat uneven because of cell motility on elongation and orientation. We can also confirm that the cells were significantly elongated and oriented in the flow direction after exposure to SS, based on the quantitative results of aspect ratio (**Fig. 6b**) and orientation angle (**Figs. 6c** and **6d**). Some differences in aspect ratio were observed between certain conditions, but there were no noticeable differences in the morphological response of the cells to the magnitude and exposure time of SS. On the other hand, the detachment rate, which quantifies cell detachment, increased significantly under 0.5 Pa conditions (**Fig. 6e**). Although there was also some increase in detachment rate under 2.3 Pa conditions, the endothelial monolayer was maintained for a longer time compared to the 0.5 Pa conditions.

**Fig. 6.**
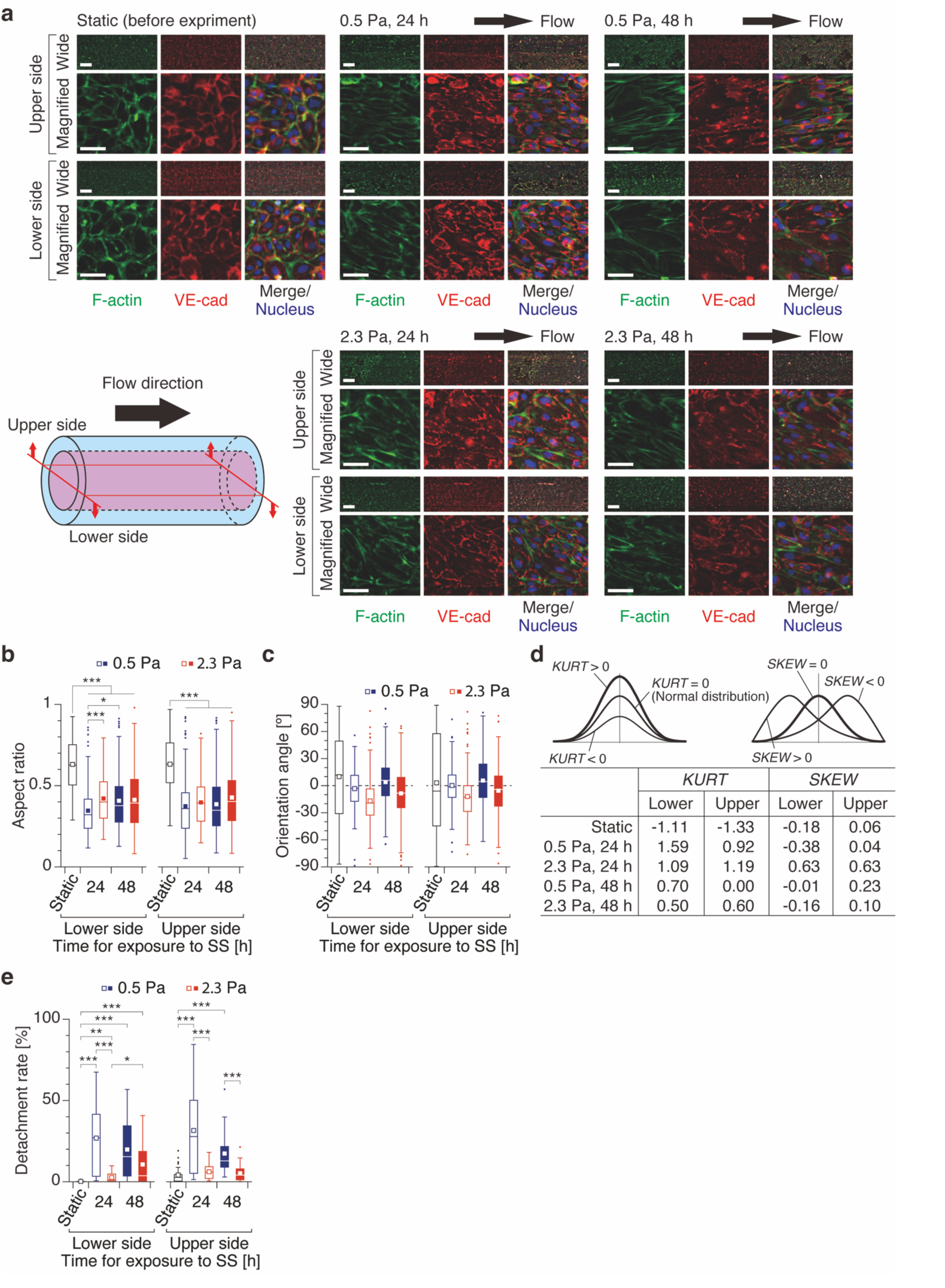
Endothelial cell responses to fluid SS in the vascular models. (**a**) Representative images of fluorescent-stained HCtAECs on the upper and lower sides of the vascular model after exposure to SS. Scale bars, 500 µm (wide view) and 50 µm (magnified view). Quantified morphological characteristics of ECs: (**b**) aspect ratio and (**c**) orientation angle. Each value was obtained from 150 (static) or 200 (SS conditions) cells. (**d**) Statistical evaluation of cell orientation distribution. Kurtosis (*KURT*) and skewness (*SKEW*) indicate the sharpness and bias of the orientation of ECs, respectively. (**e**) Cellular detachment rate on the luminal surface of the cell-cultured vascular model after exposure to SS. Each value was obtained from 30 (static) or 40 (SS conditions) images. Whiskers represent the lowest or the highest point in the data set excluding any outliers (circled dots), the box represents the 25th to 75th percentiles, the central line depicts the median, and the square inside each box indicates the average value. \**P* < 0.01, \*\**P* < 0.001, \*\*\**P* < 0.0001 (Welch’s *t* test; **b**, **c**, **e**).

### Endothelial cell dynamics after stent placement

Finally, we investigated the potential of the cell-cultured PDMS vascular model for evaluation of the endothelialization by placement of a self-expanding stent made of NiTi shape memory alloy and subsequent perfusion culture (**Fig. 7a**). Shear flow was applied 24 h after seeding of HCtAECs, and the stent was placed in the model 24 h later (**Figs. 3** and **7b**), after which the cells were further exposed to SS for up to 48 h. The shear stress level was set at 2.3 Pa. After flow exposure, the cells in the cultured vascular model with the stent in place were fixed with PFA, immunofluorescently stained, and then punctured for observation (**Fig. 7c**). ECs migrated from the lateral to the top sides of the stent at 24 to 48 h after placement (**Fig. 7d**). Expression of the intercellular adhesion protein VE-cadherin could not be confirmed. This was due to the strong reflection and scattering of fluorescence on the metal surface of the stent, and the tendency was found to be stronger for longer wavelength fluorescence (**Supplementary Fig. 3a**). By switching the fluorescence wavelength of the secondary antibody, we confirmed that VE-cadherin was expressed at the intercellular adhesion sites (**Supplementary Figs. 3b and 3c**). The endothelialization rate was quantified based on the fluorescence-stained images of F-actin to determine how much of the lateral and top sides of the stent were covered by ECs (**Fig. 7e**). On the lateral side, the average endothelialization rate was just over 20% at 24 h and increased to approximately 50% at 48 h. In contrast, the rate on the inner top side increased slightly from 24 h to 48 h, though this was less than 10%. To completely cover the inner top side with cells, further long-term perfusion culture is required. Based on the results obtained, our developed cultured vascular model has sufficient performance to be used for quantitative evaluation of stent endothelialization. To fully reproduce vascular neointimal formation, long-term perfusion culture after stent placement is necessary, whereas it is expected that the endothelial homeostasis can be sufficiently maintained by appropriate replacement of the culture medium.

**Fig. 7.**
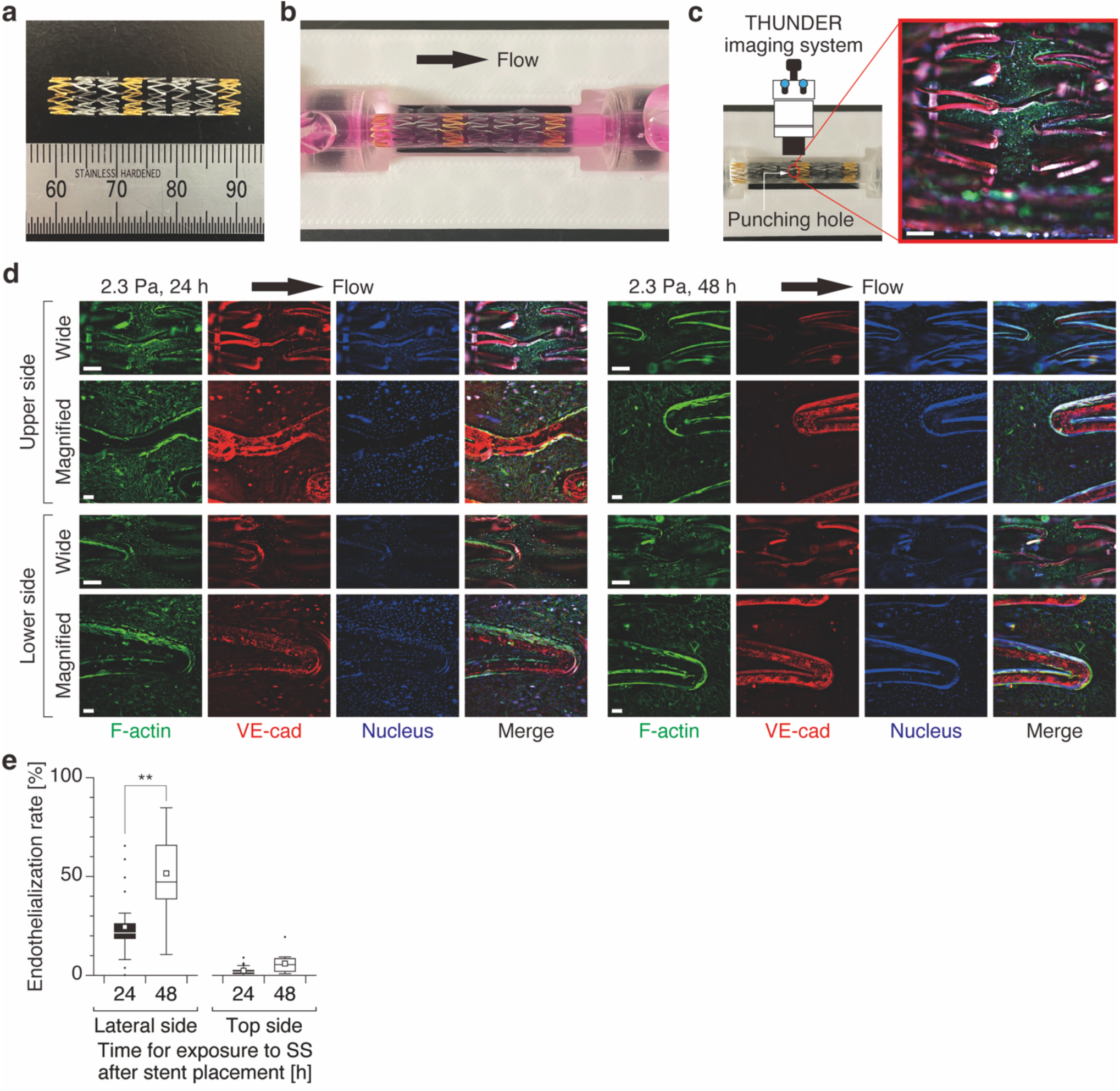
Endothelial cell dynamics in the stented vascular models exposed to SS. (**a**) A self-expanding stent used for placement in the cell-cultured vascular model. (**b**) The stent-placed vascular model. (**c**) The method to observe fluorescent-stained HCtAECs in the stented vascular model. Scale bar, 500 µm. (**d**) Representative images of fluorescent-stained ECs in the model after stent placement and exposure to SS of 2.3 Pa. Scale bars, 500 µm (wide view) and 100 µm (magnified view). (**e**) Quantitative results of endothelialization on each side of the stent placed in the cell-cultured vascular model. Each value was obtained from 25 (24-h exposure to SS) or 15 (48-h exposure to SS) locations. Whiskers represent the lowest or the highest point in the data set excluding any outliers (circled dots), the box represents the 25th to 75th percentiles, the central line depicts the median, and the square inside each box indicates the average value. \**P* < 0.01, \*\**P* < 0.001, \*\*\**P* < 0.0001 (Welch’s *t* test; **e**).

## DISCUSSION

In this study, we developed a method for constructing a cell-cultured PDMS vascular model that allows a stent to be placed, and examined its characteristics. By connecting to a flow circuit, ECs formed a monolayer covering the luminal surface of the model. They showed a morphology observed *in vivo*, *i.e.*, elongation and orientation in the direction of flow. Also, our model can reproduce the mechanical field inside the stented blood vessel, including hemodynamic stimuli. Based on the results obtained, our developed cell-cultured vascular model has sufficient performance to be used for quantitative evaluation of in-stent endothelialization. To fully reproduce vascular neointimal formation, long-term perfusion culture after stent placement is necessary, whereas it is expected that the cell system can be sufficiently maintained by appropriate replacement of the culture medium throughout the entire flow circuit.

We have successfully fabricated PDMS vascular models with a certain level of accuracy and precision, achieving a high yield (**Figs. 1** and **4**). The developed models have a wall thickness of 500 µm, which is comparable to the average thickness of medium-sized blood vessels such as carotid and coronary arteries [34,35]. However, PDMS tubes with a wall thickness of 500 µm are too stiff compared to blood vessels [36–38]. This notion is supported by the finding that the diameter change of the cell-cultured PDMS vascular model was minimal even when a stent was placed. Therefore, it is necessary to reduce the wall thickness to match the radial elasticity of the blood vessel. In the previous studies [21, 39], a cylindrical thin film was formed by dipping a core rod in PDMS or silicone resin and then rotating it to remove excess resin. Our method can also reduce the wall thickness by changing the dimensions of the core rod and outer polyacetal molds, although there might be another problem in releasing them.

Cell-cultured PDMS vascular models were constructed by seeding HCtAECs into the PDMS tubes to form their monolayers (**Figs. 2** and **5**). Shear flow generated by supplying of the culture medium is essential for maintaining endothelial monolayers, and the ECs were elongated and oriented in the direction of flow (**Fig. 6**), similar to *in vivo* conditions [40]. Hemodynamic stimuli play an important roles in mimicking the physiological state of blood vessels, while static culture can indeed be characterized as an abnormal state [41, 42]. In fact, we found that 48 h after cell seeding, endothelial cells detached from the culture surface, and the endothelial monolayer was largely disrupted. This study focused on two different magnitudes of SS, and showed that low SS (0.5 Pa) resulted in a higher detachment rate and more noticeable cell detachment than high SS (2.3 Pa) (**Fig. 6**). Human carotid artery is the source of the ECs used in the experiments, with an average SS on its wall of around from 1 to 2 Pa [43, 44]. Furthermore, given that there are reports that low SS is a risk factor for endothelial dysfunction [45], our findings regarding cell detachment at 0.5 Pa are particularly unsurprising. The cultured vascular model maintained its performance for at least 96 h after construction (cell seeding). In the case of stent placement, perfusion of the culture medium was performed from 24 h to 96 h after construction. Longer-term culture of the vascular model is technically possible, but given the degradation of EC secretions (waste products) and nutrients in the culture medium, the replacement of the medium throughout the entire flow circuit should be required at an appropriate time.

We found that it is possible to insert the stent into the cell-cultured PDMS vascular model by temporarily interrupting the perfusion culture, and this method can be used as a quantitative tool to assess the endothelialization of the implanted stent (**Fig. 7**). Of course, it should be noted that complete endothelialization of the stent requires a more prolonged perfusion culture. Some aspects of stent placement method and timing (before or during perfusion culture) were unclear in the previous studies [21, 22], but we have shown here how to do this more clearly and reproducibly (**Fig. 3**). We used a self-expanding type of stent, but the same procedure can be applied to a balloon-expandable one. The major advantage of actually placing a stent is that the mechanical field of the stented vessel can be reproduced. During stent expansion, the vascular wall is subjected to a radial force (*i.e.*, contact force) as well as a circumferential force (*i.e.*, hoop force) [6]. In the experimental setup where the stent struts are simply placed on the cell culture surface [18, 19, 20], we cannot reproduce such a complex mechanical field. For a more accurate evaluation of stent performance in an *in vitro* model, it is crucial to reproduce the mechanical field between the vessel and the stent.

We have provided the details of the method to construct a cell-cultured PDMS vascular model allowing the evaluation of in-stent endothelialization, but there are some limitations. One of the primary limitations of the cultured vascular model is that the portion of the biological tissue that exhibits vascular function is only an endothelial monolayer. Arteries are basically composed of three layers: intima, media, and adventitia. Crosstalk between SMCs in the media and ECs in the intima is important for reproducing neointimal hyperplasia, which is the cause of stenosis [46–48]. This crosstalk is also necessary for the evaluation of vascular inflammation, where the relationship between in-stent restenosis and the force exerted by the stent on the vascular wall is quantified. In the previous study, a collagen layer containing SMCs was introduced as the media [22]. The protocol to construct the cultured vascular model can be modified to include the step of placing the collagen gel inside the PDMS tube, although the culture conditions need to be optimized, and this point will be addressed in the future. Another limitation is that, as mentioned above, the vascular model has a thick and stiff wall. It should also be noted that the current thickness of 500 µm is too thick for the placing of the media (collagen layer containing SMCs) inside. Stents are actually placed so that they are embedded in the vessel wall. However, in this study, the stents did not sink into the vascular model, and changes in vessel diameter were minimal. This means that the mechanical field in the stented vessel was not accurately reproduced. To solve this problem, it is necessary to make the PDMS outer membrane thinner or to use a softer material such as alginate hydrogel instead of PDMS.

## CONCLUSION

In conclusion, this study presents a detailed method for constructing vascular models with an endothelial monolayer that allow for stent placement. By outlining step-by-step procedures for the construction of a cultured vascular model, perfusion culture, and stent placement, this work provides an essential tool not only for the *in vitro* performance evaluation (quantification of endothelialization) of stents but also for engineers involved in stent development to optimize mechanical fields in the stented blood vessel. We anticipate that this method will be a good resource to solve problems with current stent treatment, such as late restenosis and thrombosis.

## DATA AVAILABILITY

The authors declare that all data supporting the findings of this study are available within this article and its supplementary materials file, including CAD data (**Supplementary Data 2**–**17**), or from the corresponding author upon reasonable request.

## Supporting information

Supplementary Data 2-17 (CAD data)

## AUTHOR CONTRIBUTIONS

**TO:** Writing – review & editing, Visualization, Validation, Methodology, Investigation, Conceptualization. **KI:** Writing – review & editing, Validation, Methodology. **KF:** Writing – review & editing, Writing – original draft, Validation, Methodology. **DY:** Writing – review & editing, Writing – original draft, Visualization, Validation, Methodology, Supervision, Resources, Funding acquisition, Conceptualization.

## ETHICS DECLARATIONS

Ethics approval is not required. In this study, commercially available vascular ECs were used, and no animal experiments were conducted.

## SUPPLEMENTARY MATERIALS

“Cell-cultured PDMS vascular model to allow placement of implant devices” by Okuno et al.

**Supplementary Fig. 1.**
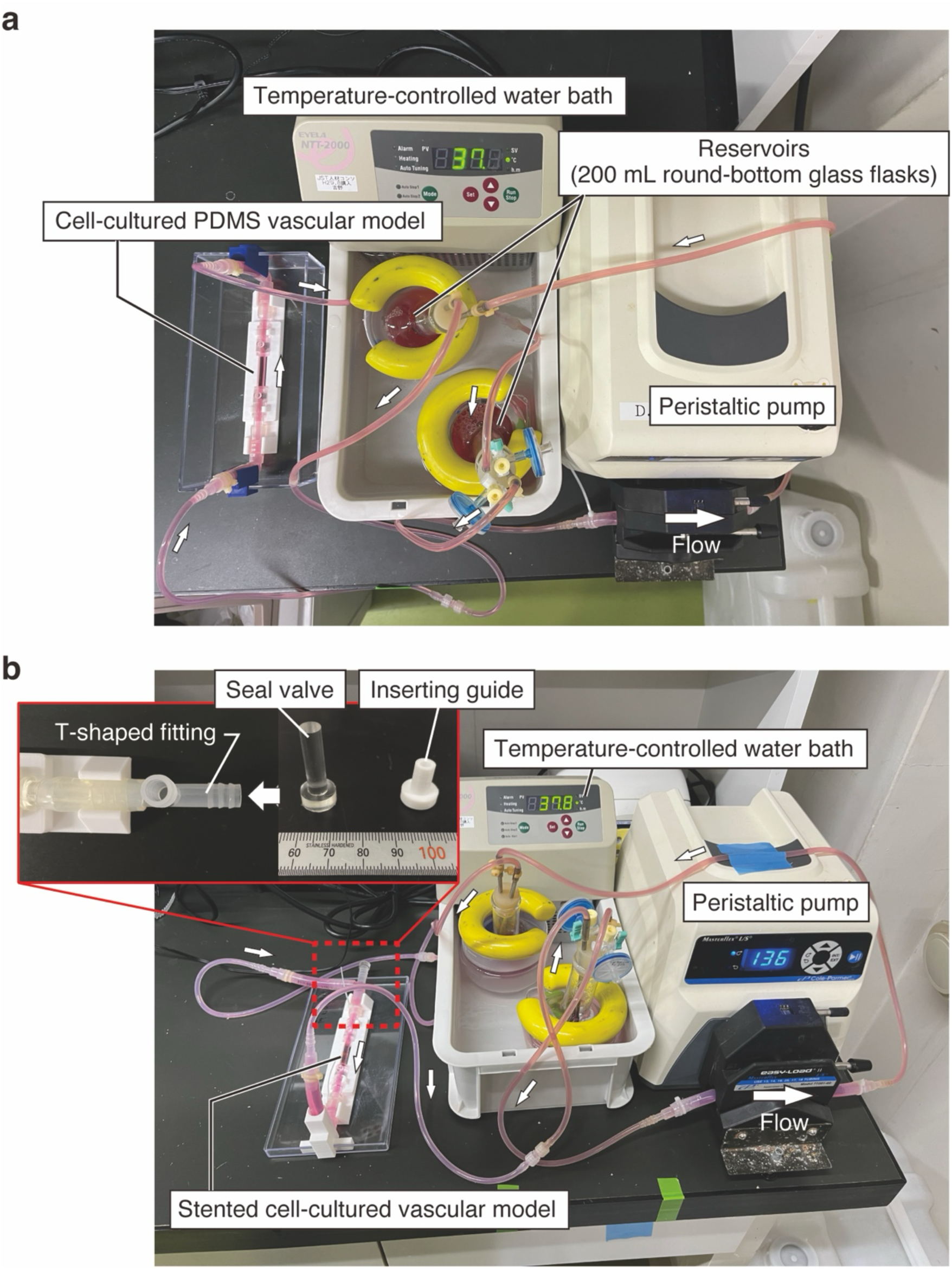
Flow-exposure cell culturing system. (**a**) Standard flow circuit and (**b**) customized one with a T-shaped fitting for stent placement experiments.

**Supplementary Fig. 2.**
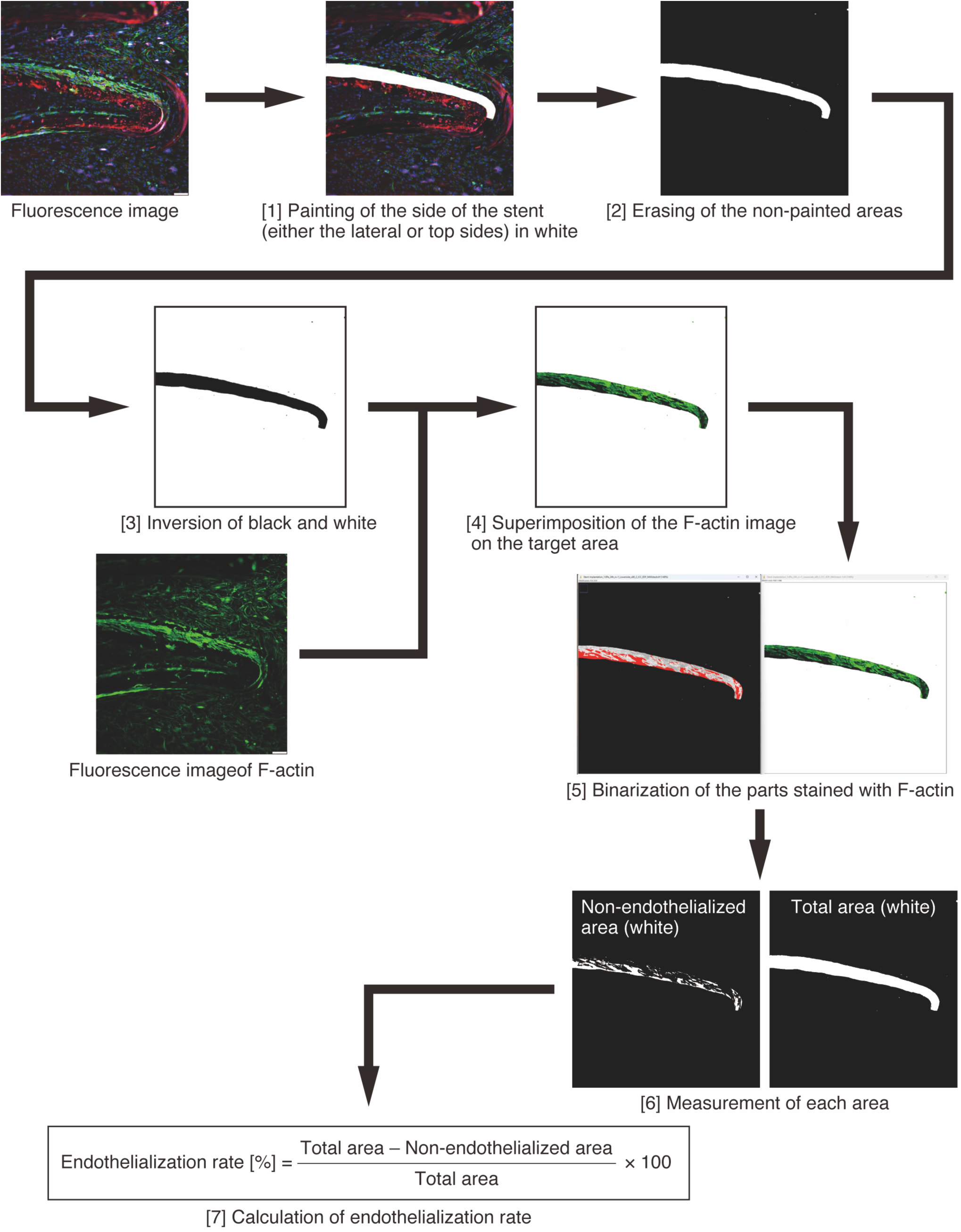
**A method for quantification of in-stent endothelialization.**

**Supplementary Fig. 3.**
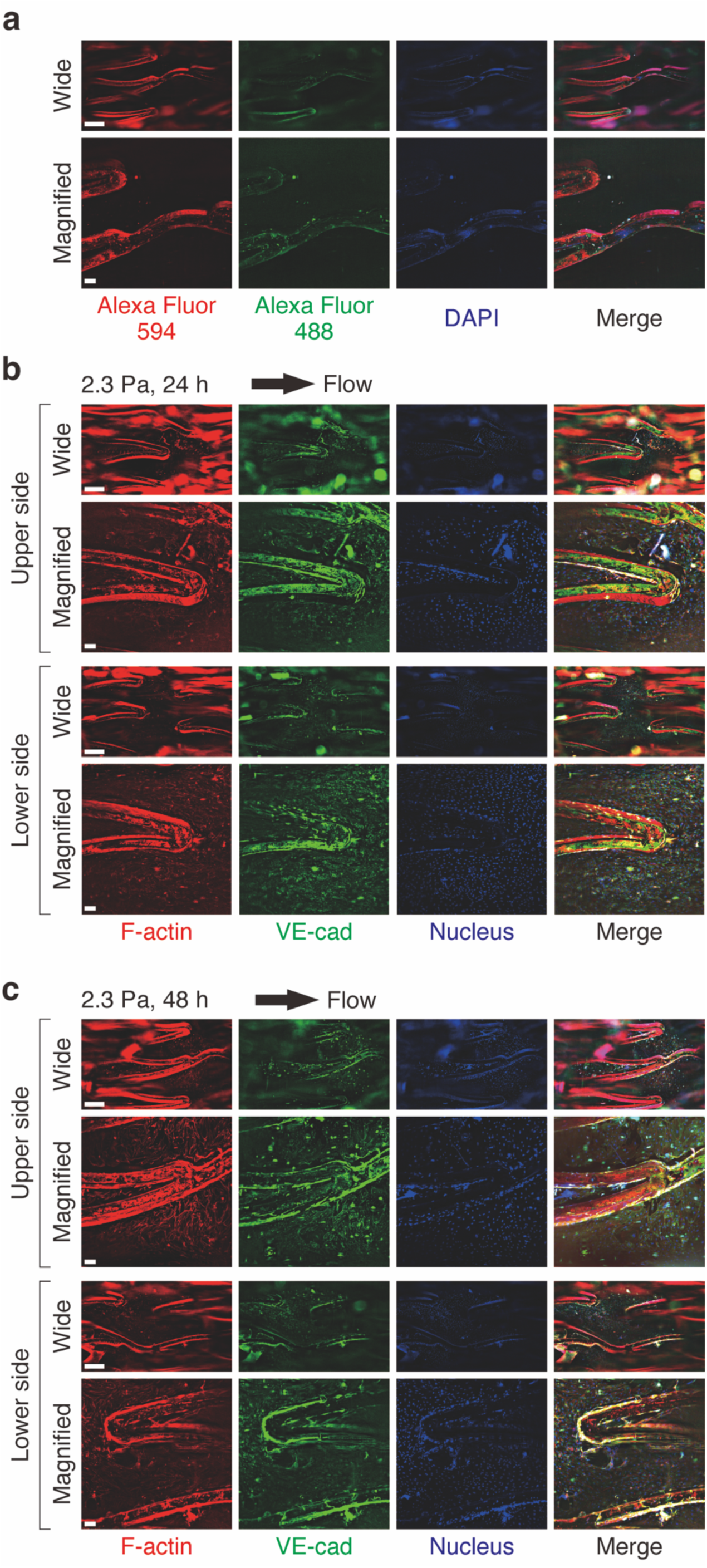
Investigation of reflection and scattering effects of fluorescence light sources. (**a**) Reflection and scattering of fluorescence on the stent surface with no cells in the vascular model. (**b**, **c**) Endothelial cell dynamics in the stented vascular models exposed to SS. Representative images of fluorescent-stained ECs in the model after stent placement and exposure to SS of 2.3 Pa for (**b**) 24 h and (**c**) 48 h. These images show the results of staining F-actin with a long-wavelength fluorescent label (594 nm) and VE-cadherin with a short-wavelength one (488 nm). Scale bars, 500 µm (wide view) and 100 µm (magnified view).

## DESCRIPTION OF ADDITIONAL SUPPLEMENTARY FILES

**File Name: Supplementary Data 1**

**Description:** The exact *P*-values and the effect size for all statistically tested data.

**File Name: Supplementary Data 2**

**Description:** Concave mold of the connectors for the inlet and outlet of the PDMS vascular model.

**File Name: Supplementary Data 3**

**Description:** Convex mold of the connectors for the inlet and outlet of the PDMS vascular model with an inner diameter of 4 mm.

**File Name: Supplementary Data 4**

**Description:** Convex mold of the connectors for the inlet and outlet of the PDMS vascular model with an inner diameter of 5 mm.

**File Name: Supplementary Data 5**

**Description:** Convex mold of the connectors for the inlet and outlet of the PDMS vascular model with an inner diameter of 6 mm.

**File Name: Supplementary Data 6**

**Description:** Mold of the PDMS vascular model with an outer diameter of 5 mm.

**File Name: Supplementary Data 7**

**Description:** Mold of the PDMS vascular model with an outer diameter of 6 mm.

**File Name: Supplementary Data 8**

**Description:** Mold of the PDMS vascular model with an outer diameter of 7 mm.

**File Name: Supplementary Data 9**

**Description:** Aluminum jig with screw holes for curing the PDMS vascular model. In the CAD data, the holes are through-ones with the pilot hole diameter, which means that it is necessary to cut an M4 tap.

**File Name: Supplementary Data 10**

**Description:** Aluminum jig with through holes for curing the PDMS vascular model.

**File Name: Supplementary Data 11**

**Description:** Mold for sealing the PDMS plugs that are used to fabricate the vascular model.

**File Name: Supplementary Data 12**

**Description:** Base jig for the cell-cultured PDMS vascular model.

**File Name: Supplementary Data 13**

**Description:** Mold for the seal valves used when seeding cells into the vascular model.

**File Name: Supplementary Data 14**

**Description:** Tapered pipe used to shrink and mount a stent into an industrial straw.

**File Name: Supplementary Data 15**

**Description:** Pushing rod used in procedures for stent placement.

**File Name: Supplementary Data 16**

**Description:** Mold for seal valves to be attached to T-fittings.

**File Name: Supplementary Data 17**

**Description:** Inserting guide for stent placement into the cell-cultured PDMS vascular model.

